# Brain networks and cognitive impairment in Parkinson’s disease

**DOI:** 10.1101/2020.12.14.422706

**Authors:** Rosaria Rucco, Anna Lardone, Marianna Liparoti, Emahnuel Troisi Lopez, Rosa De Micco, Alessandro Tessitore, Carmine Granata, Laura Mandolesi, Giuseppe Sorrentino, Pierpaolo Sorrentino

## Abstract

**Aim:** The aim of the present study is to investigate the relations between both functional connectivity and brain networks with cognitive decline, in patients with Parkinson’s disease (PD).

**Introduction:** PD phenotype is not limited to motor impairment but, rather, a wide range of non-motor disturbances can occur, cognitive impairment being one of the commonest. However, how the large-scale organization of brain activity differs in cognitively impaired patients, as opposed to cognitively preserved ones, remains poorly understood.

**Methods:** Starting from source-reconstructed resting-state magnetoencephalography data, we applied the PLM to estimate functional connectivity, globally and between brain areas, in PD patients with and without cognitive impairment (respectively PD-CI and PD-NC), as compared to healthy subjects (HS). Furthermore, using graph analysis, we characterized the alterations in brain network topology and related these, as well as the functional connectivity, to cognitive performance.

**Results:** We found reduced global and nodal PLM in several temporal (fusiform gyrus, Heschl’s gyrus and inferior temporal gyrus), parietal (postcentral gyrus), and occipital (lingual gyrus) areas within the left hemisphere, in the gamma band, in PD-CI patients, as compared to PD-NC and HS. With regard to the global topological features, PD-CI patients, as compared to HS and PD-NC patients, showed differences in multi frequencies bands (delta, alpha, gamma) in the Leaf fraction, Tree hierarchy (both higher in PD-CI) and Diameter (lower in PD-CI). Finally, we found statistically significant correlations between the MoCA test and both the Diameter in delta band and the Tree Hierarchy in the alpha band.

**Conclusion:** Our work points to specific large-scale rearrangements that occur selectively in cognitively compromised PD patients and correlated to cognitive impairment.

## Introduction

Unlike what James Parkinson claimed over two hundred years ago about the disease bearing his name, (“the senses and intellects being uninjured”) (Walshe, 1961), today we know that Parkinson’s disease (PD) is not solely a motor disease (Vitale et al., 2012). Indeed, PD is characterized by a broad spectrum of non-motor symptoms, including neuropsychiatric disturbances, autonomic dysfunctions and cognitive decline. After twenty years of disease duration, up to 80% of patients present with severe cognitive symptomatology (Aarsland et al., 2009). However, despite extensive investigation, the pathophysiological mechanisms underlying cognitive decline remain unclear (Aarsland and Kurz, 2010).

In the early stage of the disease, the brainstem and the surviving neurons of the nigrostriatal dopamine system are mostly affected by alpha synuclein depositions while, with disease progression, the neuropathological process spreads to other brain regions, including the cortex (Braak et al., 2003). Hence, PD may be regarded as a whole-brain disease.

Cognitive functions need coordinated interactions between multiple brain areas. Synchronization is one of the putative mechanisms of information routing across brain areas (Buzsáki and Draguhn, 2004). Accordingly, different electroencephalographic (EEG) or magnetoencephalographic (MEG) studies observed a relationship between neural synchrony and cognitive functions (Singer, 1999; Varela et al., 2001). Graph theory is a mathematically principled way to represent complex interactions among multiple elements. In this context, brain areas are represented as nodes, and their interactions are the links (Rubinov and Sporns, 2010; Sporns et al., 2005). Measuring topological features of the brain networks is informative about the large-scale organization underpinning cognitive processes. Recently, graph theory has been applied to MEG signals in neurodegenerative diseases, demonstrating alterations in structural organization (Pievani et al., 2014) as well as in brain functional networks, such as in amyotrophic lateral sclerosis (Sorrentino et al., 2018), hereditary spastic paraplegia (Rucco et al., 2019), and mild cognitive impairment (Jacini et al., 2018).

Given its high spatial and temporal resolution, MEG is a useful tool for detecting the evolution of brain functional connectivity. MEG systems measure the magnetic fields produced by neuronal activity, which are undistorted by the layers surrounding the brain. Therefore, it is possible to reconstruct the neural signals produced by different brain areas (source space) (Baillet, 2017). In particular, MEG has a millisecond temporal resolution, making it possible to study frequency-specific networks, and records the oscillatory activity of brain regions, allowing to estimate the phase of brain signals and, hence, synchronization (Varela et al., 2001). Typically, the canonical frequency bands (delta, theta, alpha, beta and gamma) are taken into account to understand the cognitive processes (Lopes da Silva, 2013).

Stoffers et al. have analyzed the MEG signals during resting-state in a group of *de novo* PD patients, finding changes in brain activity which included a widespread increase in theta and low alpha power, and a loss of beta and gamma power (Stoffers et al., 2007). However, they did not found correlations between brain activity and disease duration, disease stage (i.e. Hoehn and Yahr, H&Y) (Hoehn and Yahr, 1967) and disease severity (i.e. Unified Parkinson’s disease rating scale, UPDRS-III) (Fahn, 1987). The Authors hypothesized that the spectral power changes may be linked to the degeneration of non-dopaminergic ascending neurotransmitter systems. It has been demonstrated, especially in functional MRI (fMRI) studies, that the disruption of resting-state functional connectivity is important in the development of cognitive decline in PD (Amboni et al., 2018; Tessitore et al., 2012a). Some studies have compared, using MEG, the brain activity of non-demented and demented PD patients to that of matched healthy subjects. All in all, a general trend was found toward the slowing of resting brain activity in demented and (to a lesser extent) non-demented patients, as compared to healthy subjects. This slowing of oscillatory brain activity can be interpreted as a mechanism related to the progression of the disease and may be potentially involved in the development of dementia in PD (Bosboom et al., 2006; Dubbelink et al., 2013). In a source-level, resting-state MEG study, Olde Dubbelink et al. found pathologically altered functional networks in de novo PD patients (Olde Dubbelink et al., 2014) which can be interpreted as a reduction in local integration with preserved overall efficiency of the brain network. Furthermore, they have analyzed longitudinally 43 PD patients also, discovering progressive impairment in local integration in multiple frequency bands and loss of global efficiency in the PD brain network, related to a worse performance in the Cambridge Cognition Examination (CAMCOG) scale (a test assessing the global cognitive function) (Roth et al., 1986).

Ultimately, starting from the observation that the synchronization in specific frequency bands between different brain areas is the basis of a variety of cognitive processes, our hypothesis is that in PD there could be abnormal neuronal synchronization that is reflected in changes in functional connectivity and, possibly, in the topological features of the brain networks. More specifically, we hypothesize that, in PD, the progressive alteration of the brain networks would be more pronounced in patients with clinically evident cognitive impairment, as compared to cognitively unimpaired patients. To test our hypotheses, we performed a resting state MEG recording in PD patients with and without cognitive impairment, and age- and sex-matched healthy subjects (HS). We estimated synchronization between the brain source-reconstructed time series using the phase linearity measurement (PLM) (Baselice et al., 2019). We then applied the minimum spanning tree (MST) algorithm (Tewarie et al., 2015) to reconstruct the brain networks, and analyzed both functional connectivity among brain areas and topological features of the network. Finally, we correlated our results to clinical motor, cognitive and behavioral PD-specific scales.

## Materials and methods

### Participants

Thirty-nine early PD patients were diagnosed according to the modified diagnostic criteria of the UK Parkinson’s Disease Society Brain Bank (Gibb and Lees, 1988) and recruited at the Movement Disorders Unit of the First Division of Neurology at the University of Campania “Luigi Vanvitelli” (Naples, Italy). All subjects were right handed and native Italian speakers. Inclusion criteria were: a) PD onset after the age of 40 years, to exclude early onset parkinsonism; b) a modified H&Y stage ≤ 2.5. Exclusion criteria were: a) dementia associated with PD according to consensus criteria (Emre et al., 2007); b) any other neurological disorder or clinically significant or unstable medical condition; c) any contraindications to MRI or MEG recordings. Disease severity was assessed using the H&Y stages and the UPDRS III. Motor clinical assessment was performed in the “off-state” (off-medication overnight). Levodopa equivalent daily dose (LEDD) was calculated for both dopamine agonists (LEDD-DA) and dopamine agonists + L-dopa (total LEDD) (Tomlinson et al., 2010). Global cognition was assessed by means of Montreal Cognitive Assessment (MoCA) (Nasreddine et al., 2005). MoCA consists of 12 subtasks exploring the following cognitive domains: (1) memory (score range 0–5), assessed by means of delayed recall of five nouns, after two verbal presentations; (2) visuospatial abilities (score range 0–4), assessed by a clock-drawing task (3 points) and by copying of a cube (1 point); (3) executive functions (score range 0–4), assessed by means of a brief version of the Trail Making B task (1 point).

The patients were classified in two groups based on their age- and education-adjusted Italian cut-off MoCA score (Conti et al., 2015). According to these criteria we selected 20 and 19 PD patients with MoCA score respectively lower/equal (PD with cognitive impairment, PD-CI) or higher (PD with normal cognition, PD-NC) than the cut-off of 23. Depressive and apathy symptoms were assessed with the Beck Depression Index (BDI) (Beck et al., 1961) and the Apathy Evaluation Scale (AES) (Marin et al., 1991), respectively.

Twenty HS, matched for age, education and sex were also enrolled. (See Table 1).

**Table 1:** Demographic and clinical features of PD patients and healthy subjects

|  | PD-CI (n=20)<br>mean±SD | PD-NC (n=19)<br>mean±SD | HS (n=20)<br>mean±SD | p-value |
| --- | --- | --- | --- | --- |
| **Age** | 67.90±8.73 | 61.00±7.73 | 63.10±8.53 | p = 0.04 |
| **Sex (M/F)** | 10/10 | 6/13 | 11/9 | NS* |
| **Disease duration (months)** | 31.00±13.66 | 35.16±16.36 | - | NS |
| **H&Y stage** | 1.88±0.50 | 1.82±0.44 | - | NS |
| **UPDRS III** | 26.40±11.03 | 23.58±7.08 | - | NS |
| **MoCA (total)** | 19.96±2.30 | 25.05±1.63 | - | <0.001 |
| **Memory** | 0.70±0.90 | 2.32±1.49 | - | <0.001 |
| **Visuospatial abilities** | 1.75±0.99 | 3.16±0.93 | - | <0.001 |
| - **Executive functions** | 1.35±1.28 | 3.37±0.58 | - | <0.001 |
| - **Attention, concentration and working memory** | 4.35±1.49 | 5.58±0.67 | - | <0.001 |
| - **Language** | 4.30±1.23 | 5.58±0.67 | - | <0.001 |
| - **Temporal and spatial orientation** | 5.85±0.36 | 5.89±0.45 | - | NS |
| **BDI** | 5.00±5.23 | 5.37±6.87 | - | NS |
| **Apathy** | 30.25±7.14 | 29.79±6.32 | - | NS |
| **LEDD total** | 309.50±159.95 | 269.21±136.56 | - | NS |
| **LEDD DA** | 67.00±145.64 | 90.26±103.50 | - | NS |

The study was approved by the local Institutional Human Research Ethics Committee and it was conducted in accordance to the Declaration of Helsinki. All participants signed informed consent. Data are expressed as mean± standard deviation (SD). PD-CI: Parkinson’s disease patients with cognitive impairment; PD-NC: Parkinson’s disease patients without cognitive impairment; HS: healthy subjects; NS*: not significant among the three groups; NS: not significant between PD-CI and PD-NC; H&Y: Hoehn & Yahr; UPDRS: Unified Parkinson’s Disease Rating Scale; MoCA: Montreal Cognitive Assessment; BDI: Beck depression inventory; LEDD: Levodopa Equivalent Daily Dose; DA: dopamine-agonist. Note: age was statistically significant different only between PD-CI and PD-NC, with a p = 0.04.

### Magnetic Resonance Imaging acquisition

MR images were acquired on a 3-T scanner equipped with an 8-channel parallel head coil (General Electric Healthcare, Milwaukee, WI, USA) either after, or a minimum of 21 days (but not more than one month) before the MEG recording. Three-dimensional T1-weighted images (gradient-echo sequence Inversion Recovery prepared Fast Spoiled Gradient Recalled-echo, time repetition = 6988 ms, TI = 1100 ms, TE = 3.9 ms, flip angle = 10, voxel size = 1 x 1 x 1.2 mm3) were acquired.

### MEG acquisition

The MEG system acquires the signals of 163 magnetometers placed in a magnetically shielded room (AtB Biomag, Ulm, Germany). Specifically, 154 sensors cover the entire head of the subject; the remaining ones, organized into three orthogonal triplets, are positioned more distant from the helmet and used to measure and reduce the environmental noise (Lardone et al., 2018; Sorrentino et al., 2017). MEG data were acquired during two, eyes-closed, resting state segments, each 3.5 minutes long. The patients were in the off-state (i.e. after drug withdrawal for 24 hours, without the effects of the therapy).

In order to reconstruct the position of the head in the helmet during the MEG, we digitalized, before acquisition, the position of four reference coils (attached to the head of the subject) and four anatomical landmarks (nasion, right and left pre-auricular and apex) using Fastrak (Polhemus^®^). The coils were activated before each segment of the registration. During the MEG acquisition, electrocardiographic (ECG) and electrooculographic (EOG) signals were also recorded to remove physiological artefact (Gross et al., 2013; Rucco et al., 2019). After an anti-aliasing filter, the data were sampled at 1024 Hz.

### Preprocessing

The MEG data were filtered in the band 0.5-48 Hz using a 4th-order Butterworth IIR band-pass filter, implemented offline using Matlab scripts within the Fieldtrip toolbox (Oostenveld et al., 2011). To reduce the environmental noise, Principal Component Analysis (PCA) was used (de Cheveigné and Simon, 2007; P.K. Sadasivan, 1996). Subsequently, an experienced rater identified the noisy channel/segments of acquisition through visual inspection. On average, 140 ± 4 channels were used. After that, Independent Component Analysis (ICA) (Barbati et al., 2004) was performed to identify and remove ECG (typically 1-2 two components) and EOG (0-1 components) signals from the MEG data.

### Source reconstruction

The subject’s anatomical landmarks were visually identified on the native MRI of the subjects and used to co-register the MEG acquisition, which was then spatially normalized to a template MRI. Subsequently, the time series related to the centroids of 116 regions-of-interest (ROIs), derived by the Automated Anatomical Labelling (AAL) atlas (Gong et al., 2009; Tzourio-Mazoyer et al., 2002) were reconstructed based on Nolte’s volume conduction model (Nolte, 2003) and the Linearly Constrained Minimum Variance (LCMV) beamformer algorithm (Van Veen et al., 1997). However, we considered only the first 90 ROIs, excluding those representing the cerebellum, given the low reliability of the reconstructed signal in those areas. For each ROI, we projected the time series along the dipole direction that explained most variance by means of singular value decomposition (SVD), using Fieldtrip toolbox (Oostenveld et al., 2011).

The beamformer estimates the temporal series representing the activity of the brain regions. Such signals are filtered in the five canonical frequency bands (delta (0.5 – 4 Hz), theta (4.0 – 8.0 Hz), alpha (8.0 – 13.0 Hz), beta (13.0 – 30.0 Hz) and gamma (30.0 – 48.0 Hz)), and analysed separately.

### Connectivity analysis

To evaluate the synchronization between brain regions, we adopted the Phase Linearity Measurement (PLM) (Baselice et al., 2019). This novel, undirected metric, developed by our group, measures the synchronization between brain regions exploiting the power spectrum of their phase differences in time. It is defined as follows:

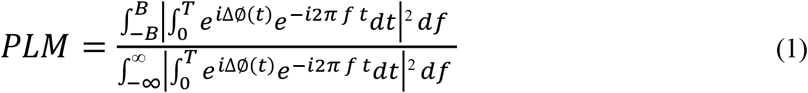

where the 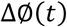 represent the phase difference between two signals, 2B is the integration band, *f* is the frequency and *T* is the observation time interval. The PLM ranges between 0 and 1, where 1 indicates perfect synchronization and 0 indicates non synchronous activity.

Based on PLM, we obtained a 90×90 weighted adjacency matrix for each temporal series (with a duration > 4s), for each subject, in each frequency band.

Starting from these weighted adjacency matrices we calculated, for each ROI, the nodal PLM for each ROI as the average PLM between a specific ROI and all other ROIs, and the global PLM as the average of all nodal PLM values.

### Network analysis

Starting from the weighted adjacency matrices, we reconstructed, based on the minimum spanning tree (MST) algorithm, a binary network, where the 90 areas of the AAL atlas are the nodes and the entries represent the edges.

To describe the network, we computed nodal centrality measures (degree, betweenness centrality) and global, non-centrality (leaf fraction, degree divergence, diameter, tree hierarchy) metrics (Stam et al., 2014; Tewarie et al., 2015). The degree of a node is defined as the number of links incident on a given node. The betweenness centrality (BC) is the number of shortest paths passing through a given node over all the shortest paths of the network (Freeman, 1977). The leaf fraction (Lf) is the fraction of leaf nodes in the MST, where a leaf node is defined as a node with degree one (Boersma et al., 2013). The degree divergence (K) measures the broadness of the degree distribution (Tewarie et al., 2015). The diameter is defined as the longest shortest path of the MST. Lastly, the tree hierarchy (Th) is the number of leaves over the maximum betweenness centrality.

### Statistical analysis

To test differences in age and sex among the three groups we use ANOVA and the Chi square, respectively, after checking the normal distribution of variables. Clinical parameters, between PD-CI and PD-NC patients, were compared using t-test.

The three groups were compared for each variable of interest (connectivity and topological metrics) using the permutational analysis of variance (PERMANOVA), a non-parametric test in order to evaluate the effect of cognitive impairment on brain connectivity in PD-CI, PD-NC patients and in controls. Then, all the p-values were corrected using the false discovery rate (FDR) (Benjamini and Hochberg, 1995), so as to account for multiple comparison between the variables. For the significant p values (after FDR correction), post-hoc analysis was carried out, using Scheffe’s correction for multiple comparisons among groups.

To correlate the connectivity and topological metrics with the clinical scales, we used the Spearman’s rank correlation coefficient. All statistical analyses were performed using custom scripts written in Matlab 2018a. The significance level was set at p < 0.05.

## Results

### Population Characteristics

The studied population consists of 20 PD-CI, 19 PD-NC patients and 20 HS. The gender among the three groups showed no significant difference. PD-NC were slightly younger than PD-CI patients (p = 0.04), while no difference were found in terms of disease duration, disease stage (i.e. H&Y stage), motor impairment (i.e. UPDRS III), depression (i.e. BDI scale) and apathy (i.e. AES) between the two PD subgroups. As expected, significant differences were found in terms of MoCA scale and its subtests between PD-CI and PD-NC patients (Table 1).

### MEG data

#### Connectivity analysis

Regarding the global PLM value, we found a statistical significant difference in the gamma band among the groups with a p = 0.0416 (H (2,58) = 3.365), with post-hoc analysis showing that PD-CI patients differed from HS, having lower global PLM, see Fig. 1.

**Fig. 1.**
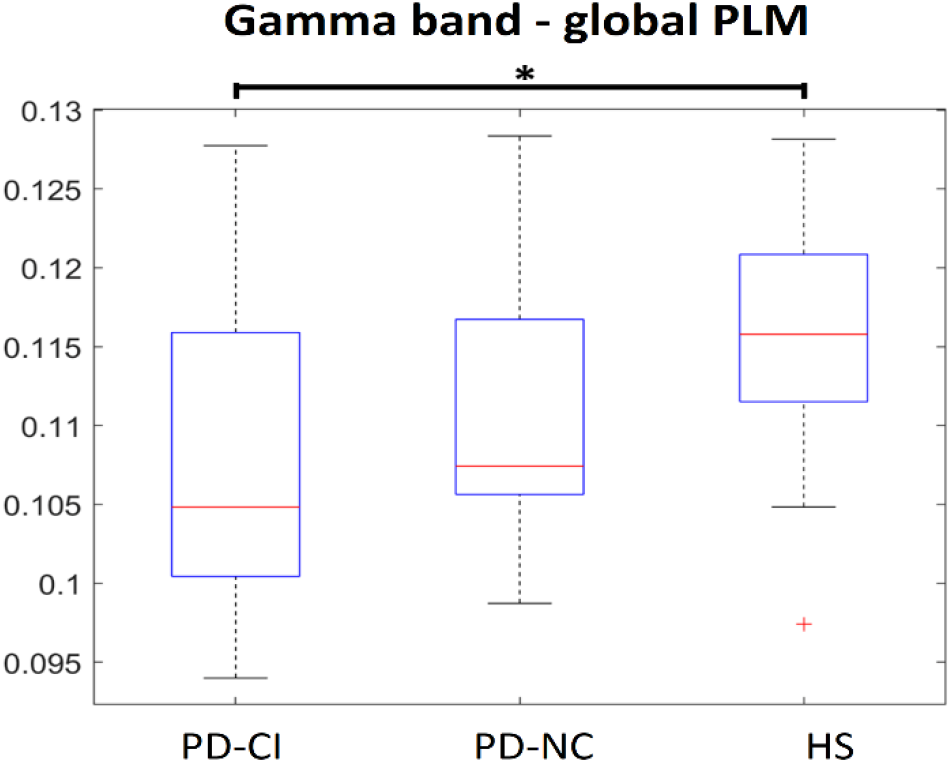
Differences in the global PLM value among PD-CI, PD-NC and HS. The box plots refer to differences in the global PLM value in gamma band among PD-CI, PD-NC and HS. The upper and lower bound of the box refer to the 25^th^ to 75^th^ percentiles, the median value is represented by horizontal line inside each box, the whiskers extent to the 10^th^ and 90^th^ percentiles, and further data are considered as outliers and represented by the symbol +. PD-CI group shows a lower global PLM value as compared to both PD-NC group (without reaching statistical significance) and HS (* = p < 0.05).

When we compared the nodal PLM values among the three groups, we found differences in the gamma band in the following areas of the left hemisphere: Postcentral gyrus (H (2,58) = 6.578, p = 0.002, pFDR = 0.039), Lingual gyrus (H (2,58) = 7.563, p = 0.001, pFDR = 0.039), Fusiform gyrus (H (2,58) = 9.279, p < 0.001, pFDR = 0.036), Heschl’s gyrus (H (2,58) = 6.985, p = 0.002, pFDR = 0.039), inferior Temporal gyrus (H (2,58) = 7.377, p = 0.001, pFDR = 0.039). In the post-hoc analysis, PD-CI patients showed a lower PLM value with respect to HS in all significant ROI, while PD-NC patients only reached statistical significance in the left lingual and the left Fusiform areas, as showed in Fig. 2.

**Fig. 2.**
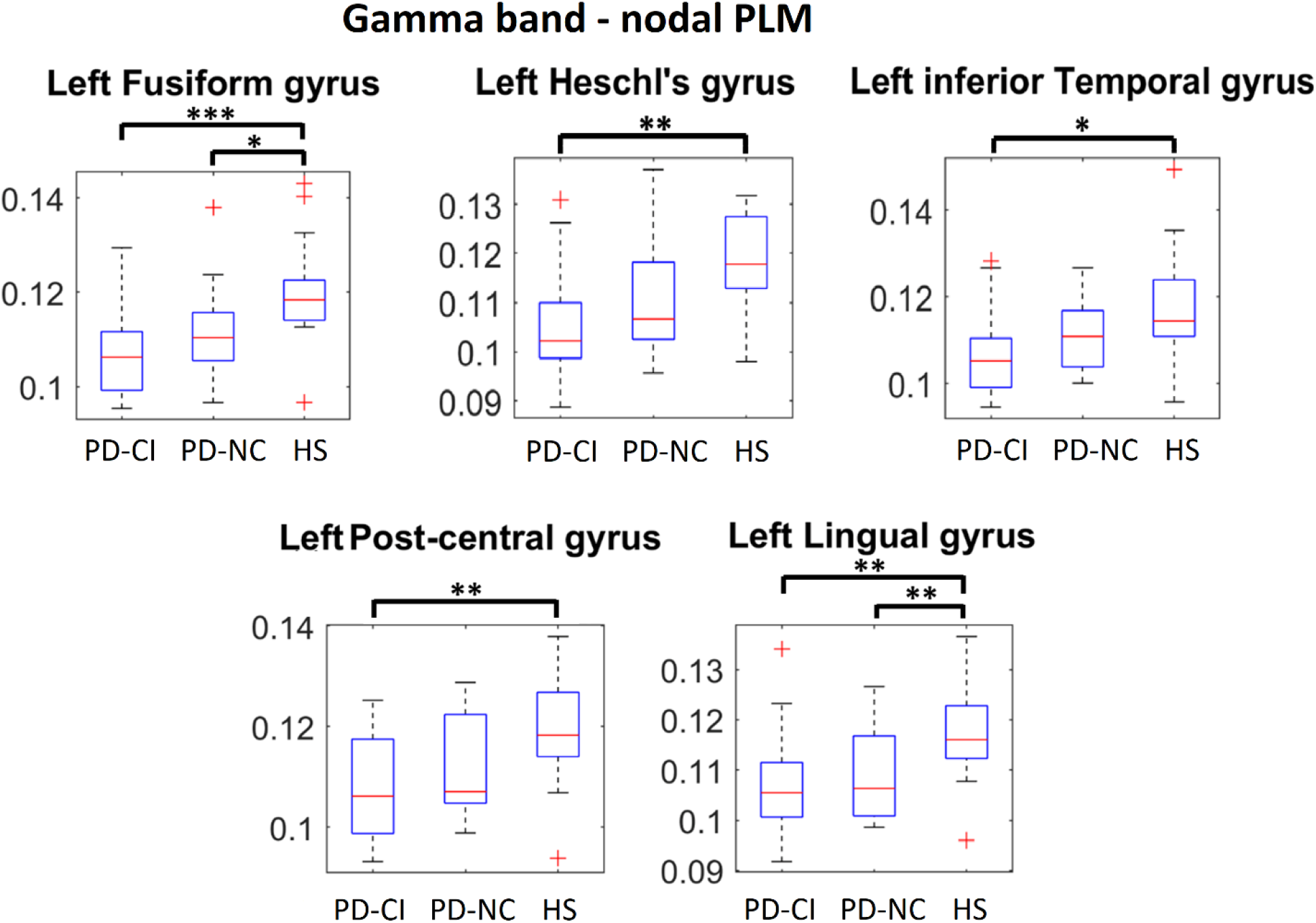
Differences in the nodal PLM values among PD-CI, PD-NC and HS. The box plots refer to differences in the nodal PLM value in gamma band among PD-CI, PD-NC and HS. The upper and lower bound of the box refer to the 25^th^ to 75^th^ percentiles, the median value is represented by horizontal line inside each box, the whiskers extent to the 10^th^ and 90^th^ percentiles, and further data are considered as outliers and represented by the symbol +. PD-CI group shows lower nodal PLM values in Fusiform gyrus, Heschl’s gyrus and Inferior temporal gyrus, Post-central gyrus, Lingual gyrus, on the left, as compared to both PD-NC group and HS. * = p < 0.05, ** = p < 0.01, *** = p < 0.001

#### Topological network analysis

We found topological differences in the brain networks among PD-CI, PD-NC and HS, in different frequency bands. With respect to Lf, differences appeared in the delta (H (2,58) = 4.732, p = 0.012, pFDR = 0.049), the alpha (H (2,58) = 4.371, p = 0.017, pFDR = 0.028) and the gamma band (H (2,58) = 7.052, p = 0.002, pFDR = 0.012). Post-hoc analysis showed that, in all the three bands, PD-CI patients had higher leaf fraction as compared to HS, as depicted in Fig. 3.

**Fig. 3.**
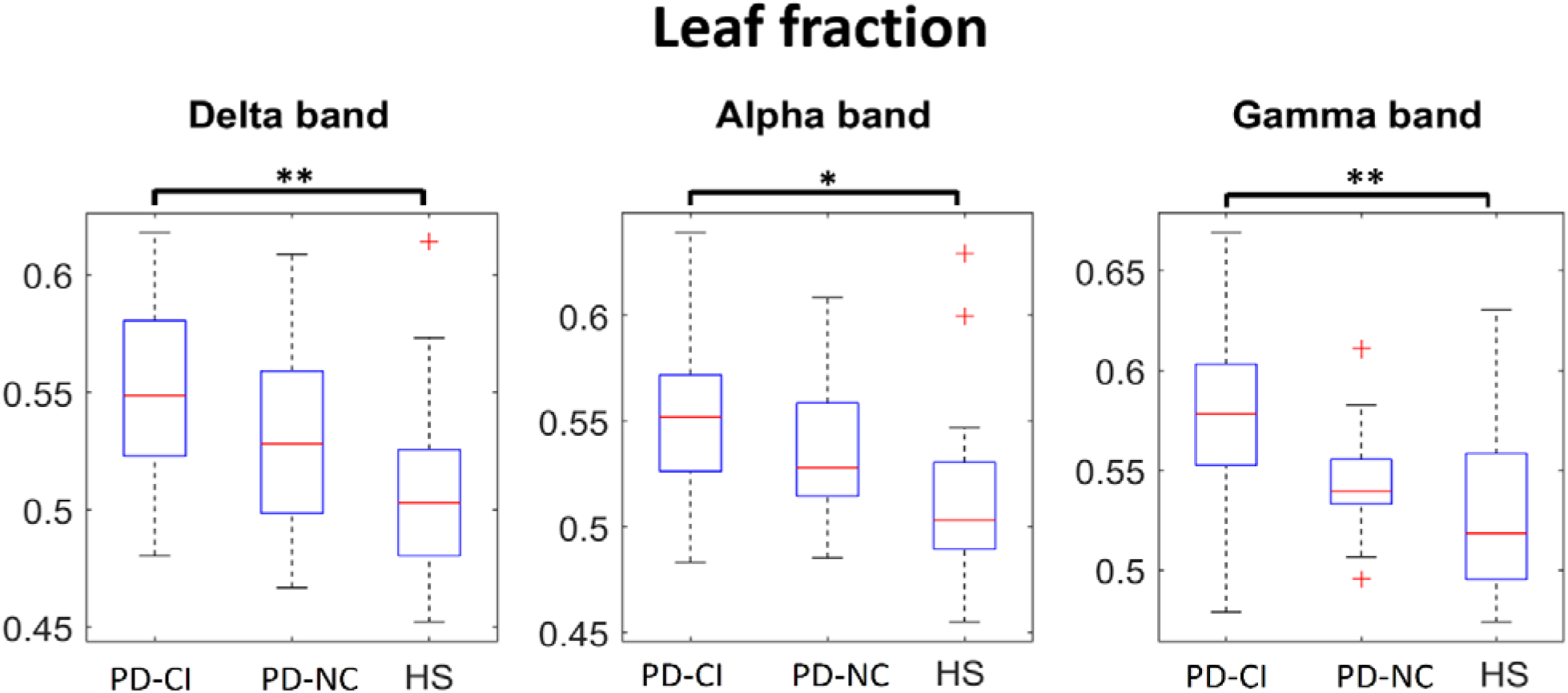
Differences in Leaf fraction parameter, among PD-CI, PD-NC and HS. The box plots refer to differences in the Lf among respectively PD-CI, PD-NC and HS. The upper and lower bound of the box refer to the 25^th^ to 75^th^ percentiles, the median value is represented by horizontal line inside each box, the whiskers extent to the 10^th^ and 90^th^ percentiles, and further data are considered as outliers and represented by the symbol +. PD-CI group shows a higher Lf, compared to both PD-NC group and HS, in delta, alpha and gamma band. * = p<0.05, ** = p<0.01

The Th differed among the three groups in the alpha (H (2,58) = 5.329, p = 0.006, pFDR = 0.016) and the gamma band (H (2,58) = 5.523, p = 0.007, pFDR = 0.019). In the post-hoc analysis, both PD-CI and PD-NC patients differed from HS with a higher Th in the alpha band, but only PD-CI patients differed from the HS in the gamma band, as reported in Fig. 4.

**Fig. 4.**
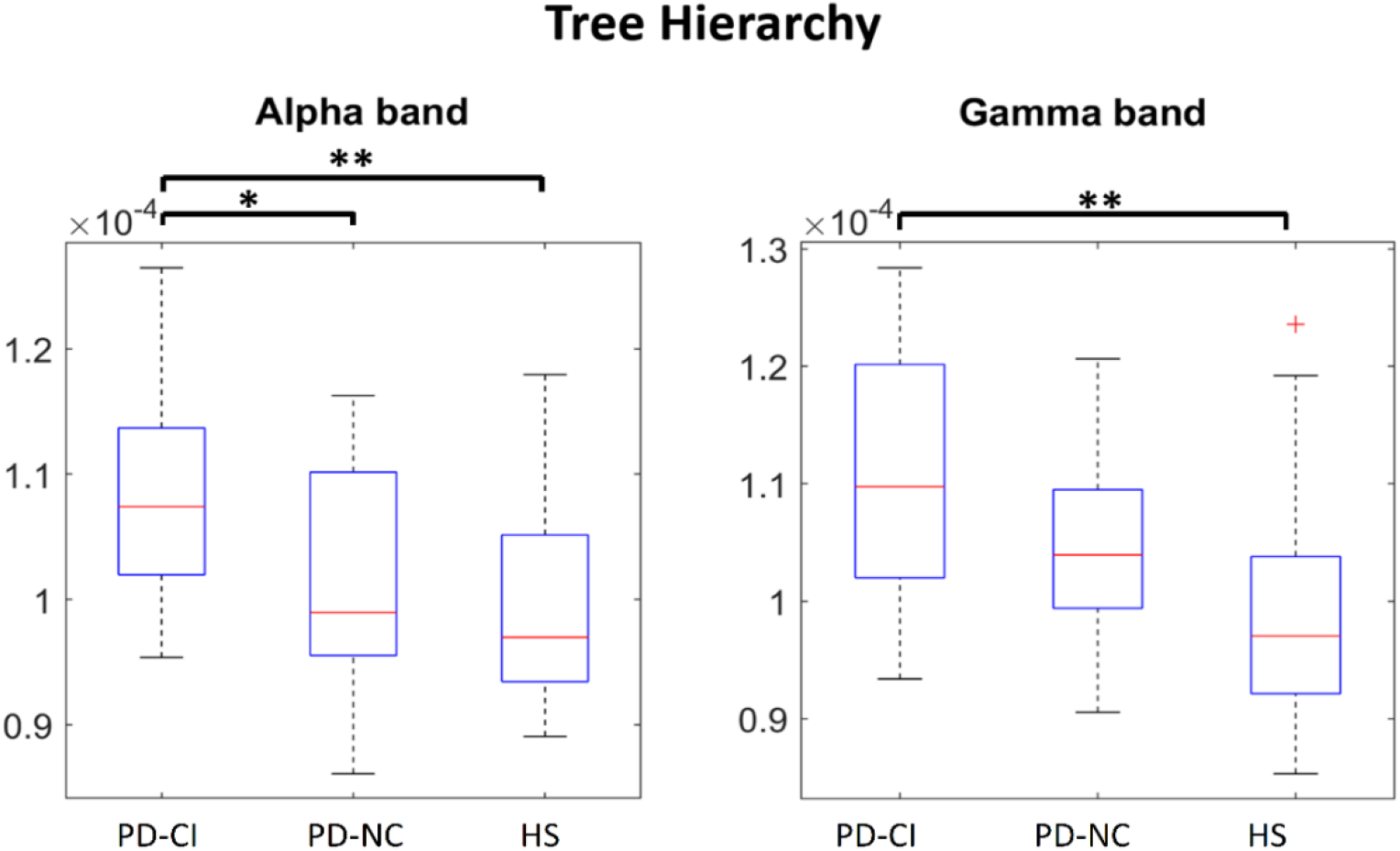
Differences in Tree Hierarchy parameter among PD-CI, PD-NC and HS. The box plots refer to differences in the Th among respectively PD-CI, PD-NC and HS. The upper and lower bound of the box refer to the 25^th^ to 75^th^ percentiles, the median value is represented by horizontal line inside each box, the whiskers extent to the 10^th^ and 90^th^ percentiles, and further data are considered as outliers and represented by the symbol +. The PD-CI group shows a higher Th, compared to both PD-NC group and HS, in the alpha and gamma bands. * = p<0.05, ** = p<0.01

The diameter was statistically different in the delta band (H (2,58) = 4.214, p = 0.019, pFDR = 0.049) among the three groups, and in particular between PD-CI patients and HS, see Fig. 5.

**Fig. 5.**
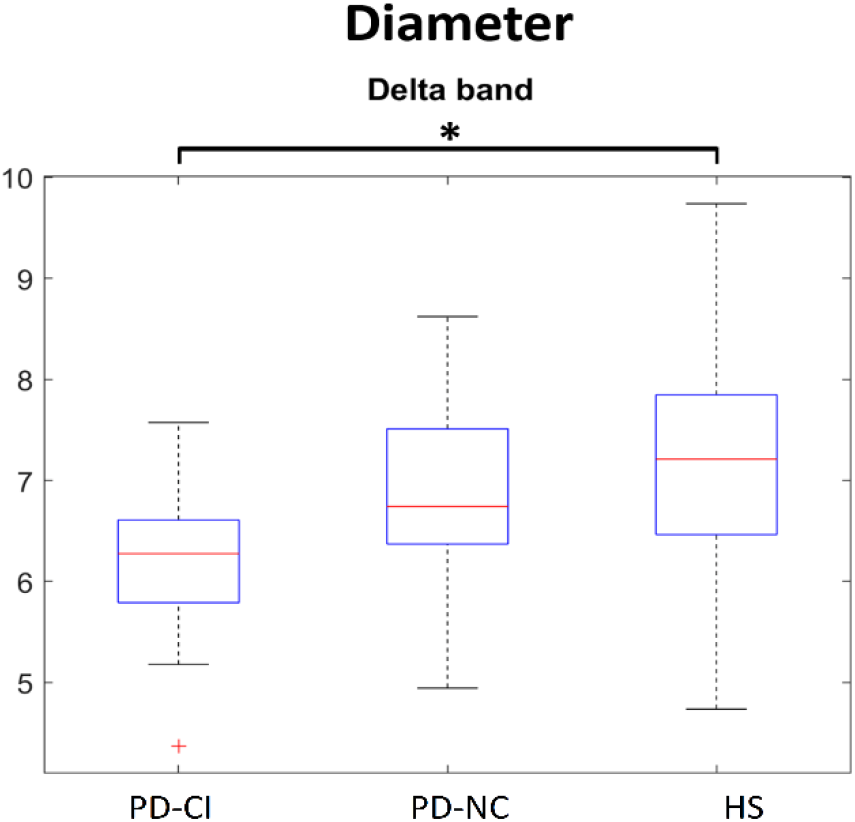
Differences in Diameter in PD-CI, PD-NC and HS. The box plots refer to differences in the D among respectively PD-CI, PD-NC and HS. The upper and lower bound of the box refer to the 25^th^ to 75^th^ percentiles, the median value is represented by horizontal line inside each box, the whiskers extent to the 10^th^ and 90^th^ percentiles, and further data are considered as outliers and represented by the symbol +. PD-CI group shows a statistically significant lower Diameter compared to both PD-NC group and HS, in delta band. * = p<0.05

However, it is to be noted that, although most of the parameters in the PD-NC group did not reach statistical significance, a trend seems evident nonetheless, such that cognitively unimpaired patients show intermediate values between healthy controls and cognitively compromised patients. No statically significant difference was found among the three groups in the K, the other global topological parameters calculated, and in the centrality parameters.

#### Correlations analysis

As shown in Fig. 6, we found a statistically significant correlation between the MoCA total score and both the Diameter in delta band (R = 0.352, p = 0.028), and the Tree Hierarchy in the alpha band (R = −0.374, p = 0.019). No other statistically significant correlation between connectivity metrics and clinical scales was found.

**Fig. 6.**
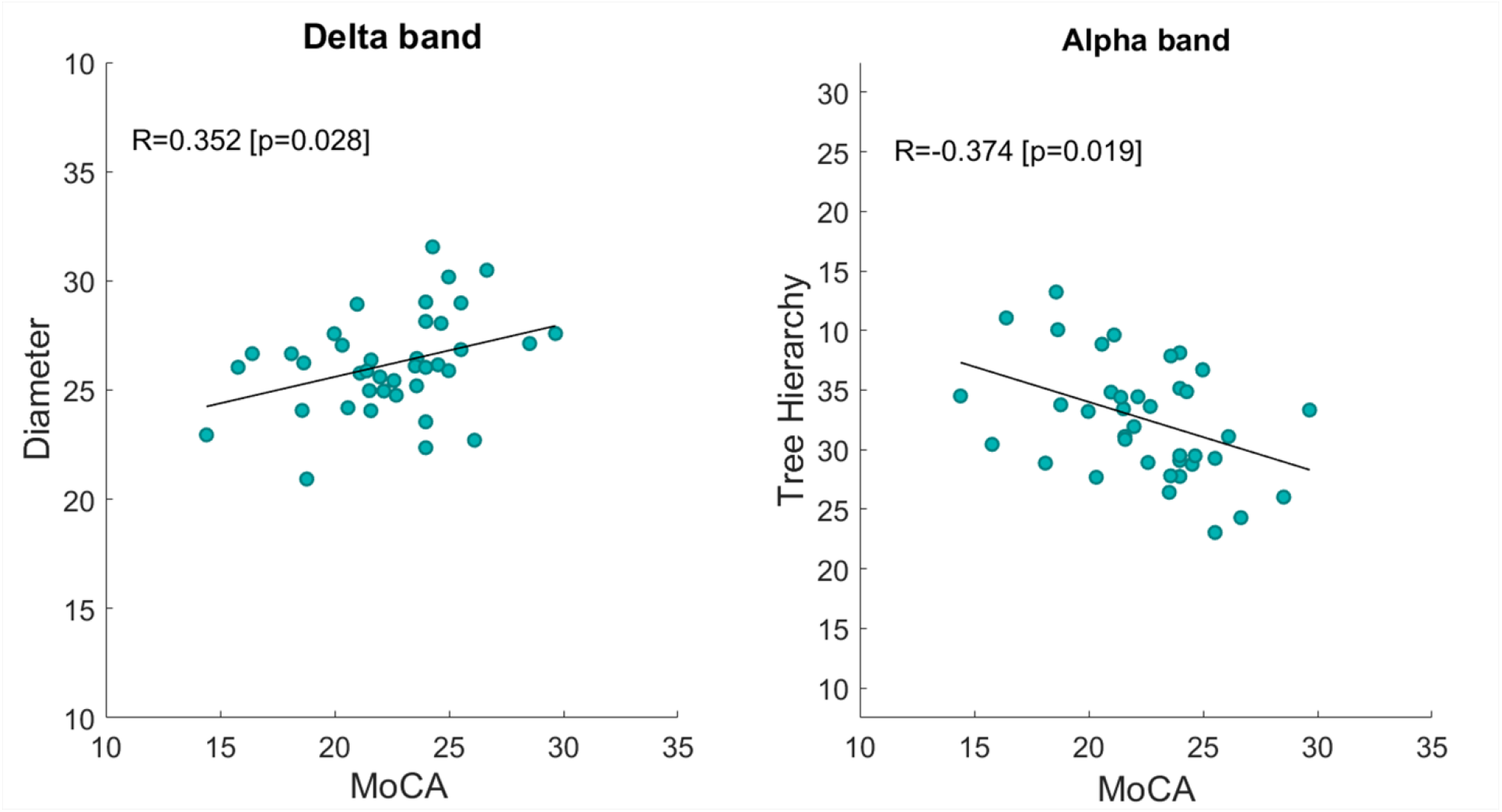
Spearman’s rank correlation coefficient. MoCa test correlates positively with the Diameter (R = 0.352, p = 0.028) and negatively with the Tree Hierarchy (R = 0.374, p = 0.019).

## Discussion

Our study was designed to test the hypothesis that the cognitive decline observed in PD patients may be associated to specific changes of both functional connectivity and brain topology. Furthermore, we hypothesized that the extent of brain network alterations may be correlated with the cognitive outcome. By applying the PLM, a connectivity metric that measures the synchronization between brain regions, (Baselice et al., 2019) to MEG signals, we were able to highlight differences in the global and nodal PLM values in PD-CI as compared to both PD-NC and HS. Furthermore, using graph analysis, we found specific PD-related changes in brain network topology which were related to cognitive functioning.

### Functional connectivity

We found that the global PLM value in the gamma band was significantly reduced in PD-CI patients as compared to HS. This measure, obtained by averaging over all 90 (one for each ROI) nodal PLM values, is a measure of global functional connectivity. Interestingly, the global PLM of PD-NC patients was intermediate between that of HS and PD-CI (although the difference was not statistically significant).

The nodal PLM values showed a similar trend to that of the global PLM. For example, the nodal PLM of cognitively PD-NC patients was intermediate between PD-CI patients and HS in the gamma band. Specifically, a statically significant reduction of the functional connectivity was observed in several temporal (fusiform gyrus, Heschl’s gyrus and inferior temporal gyrus), parietal (postcentral gyrus), and occipital (lingual gyrus) areas within the left hemisphere, as compared to HS. Moreover, the PLM of the lingual and fusiform left gyri was significantly reduced with respect to the HS in both PD-CI and PD-NC patients (Fig. 2).

The heterogeneity of the clinical onset, the prognostic evolution as well as the response to dopaminergic therapy suggest the existence of two distinct cognitive syndromes in PD (although with overlapping elements), namely the frontostriatal syndrome (Tessitore et al., 2012b) and the posterior cortical syndrome (Baggio et al., 2015; Tremblay et al., 2013). The former is cognitively characterized mainly by dysexecutive disorders, and is strictly related to the dopaminergic imbalance (Gotham et al., 1986), while in the latter, memory deficit, visuospatial/visuoperceptual disturbances and more generally global cognitive decline are frequently observed (Williams-Gray et al., 2009). Importantly, the posterior cortical syndrome is associated with a worse cognitive prognosis (Kehagia et al., 2010). Overall, our results are in line with this view, where the form presenting the greater risk of developing dementia (Olde Dubbelink et al., 2014) showed widespread functional connectivity in temporal, parietal and occipital regions (Baggio et al., 2015). Interestingly, cortical areas showing reduced synchronization in cognitively impaired PD subjects (i.e. fusiform gyrus, Heschl’s gyrus, inferior temporal gyrus, postcentral gyrus, and lingual gyrus) are mainly involved in the posterior cortical syndrome.

Taking into account the clinical evidence suggesting that damage in such regions leads to severe cognitive impairment with a high risk of developing dementia, we might speculate that, if these regions are less integrated with the rest of the brain, then the cognitive functioning might be impaired. This is also supported by our correlation analysis, showing that the less is the synchronization between these areas and the rest of the brain, the worst the cognitive performance. It is important to note that there was a clear downward trend between HS and all PD in both global and nodal PLM values, with PD-NC group always displaying intermediate values. This observation could suggest that the reduction of the functional connectivity in terms of reduced overall synchronization (estimated by the PLM) progresses till to exceed a threshold, the cognitive impairment acquires clinical significance (Sorrentino et al., 2020). It is even more interesting to observe that the reduction in synchronization in the posterior regions (along with the cognitive impairment), is not a function of disease progression or severity, as documented by the comparison of the clinical scales between the two PD groups.

Finally, it is worth noting that all these results are in the gamma band (30-48 Hz), which has been related to visual perception, attention, auditory processing, learning and memory (Hoogenboom et al., 2006; Kaiser and Lutzenberger, 2005). Interestingly, dopamine agonists have been shown to increase gamma-band activity in both cortical and subcortical networks (Brown, 2003).

### Brain network topology

The reduction of functional connectivity in PD patients is linked to changes in the large-scale functional organization of the brain, as captured by our topological network results. With regard to the centrality parameters (degree and betweenness centrality), which evaluate the topological characteristic of each single region, we did not find any statistically significant difference among the three groups. However, with regard to the global parameters, expressing global topological features of the brain network, PD-CI patients, as compared to HS and PD-NC patients, showed widespread differences in multi frequencies bands (delta, alpha, gamma) in the Lf, Th (both higher in PD-CI) and Diameter (lower in PD-CI) (Fig. 3, 4 and 5).

It should be noted that, similarly to the functional connectivity, PD-NC group shows an intermediate profile between HS and PD-CI, even when the difference does not reach statistical significance (see Fig. 4).

The Lf is defined as the ratio between the number of leaf nodes (nodes with degree = 1) and the maximum possible number of links (total number of nodes minus 1). A Lf equal to 1 indicates a network with a star-like topology (Tewarie et al., 2015), where each couple of nodes is topologically closer, and most shortest path pass on a small subset of highly-important nodes. On the contrary, a Lf equal to 0 signifies a line-like network, which is less reliant on any singly node, and hence with higher resiliency to targeted attacks (Rubinov and Sporns, 2010; Tononi et al., 1994). Related to the Lf, the Diameter provides information about the distance between all pairs of nodes. In fact, lower Diameter, as showed by PD patients in the delta band, is indicating a more compact, star-like network (Boersma et al., 2013). Finally, the tree hierarchy quantifies the trade-off between efficient communication (large-scale integration) and prevention of the overload of the most important nodes. A higher tree hierarchy, as found in PD-CI, may suggests a sub-optimal balance, with respect to both PD-NC (in the alpha band) and HS (in the alpha and the gamma band), in the sense that, in pathology, the network integration becomes reliant on a small subset of important areas, hence losing resiliency. This mechanism might underlie the reduction of functional connectivity found in some brain areas (see Fig 1 and 2) linked to cognitive deterioration.

### Correlation analysis

Interestingly, as reported in Fig. 7, we found statistically significant correlation between the MoCA test and both the Diameter in delta band (direct correlation) and the Tree Hierarchy in the alpha band (inverse correlation). These correlations are in line with our findings and support the hypothesis of reduced synchronization in some brain areas, as well as hyperconnected network topology, that might capture sub-optimal large-scale functional organization underpinning cognitive impairment development in PD patients.

## Conclusion

In conclusion, in this work, we show that in PD patients in the early phase of the disease, the functional connectivity changes, as well as the topological rearrangements within the large-scale functional networks, are correlated to cognitive impairment. In particular, we found reduced functional connectivity in PD-CI (with respect to both PD-NC and HS) in terms of reduced overall synchronization, as estimated by the PLM, as well as specifically in the posterior hubs. Furthermore, analyzing the brain networks, we found a more star-like topology in PD-CI.

It is noteworthy to observe that both PD groups (i.e. PD-CI and the PD-NC group) did not differ with regard to the disease stage as well as to the motor impairment. Nonetheless, the group affected by earlier development of cognitive impairment was the one showing reduced synchronization in the posterior areas. These data are in line with the hypothesis that two distinct clinical phenotypes (although with overlapping elements) exist and that involvement of the posterior regions relates to earlier cognitive decline.

## Author contributions

RR collected and acquired the dataset, processed the data and conceptualized the study; ML, ETL and FB processed the data; AL, RDM and AT collected the sample; CG, LM, GS contributed to interpreting the results and critically revised the article; PS supervised the study. All authors interpreted the results and wrote the manuscript.

## Data availability

The MEG data and the reconstructed avalanches are available upon reasonable request to the corresponding author, conditional on appropriate ethics approval at the local site.

